# Mutational Landscape of TP53 Across Cancer Types

**DOI:** 10.1101/2025.08.12.669884

**Authors:** Maria Abran

## Abstract

The *TP53* gene is the most frequently mutated tumor suppressor in human cancers, but its mutation patterns vary across tumor types. Using data from *cBioPortal* (downloaded 9 August 2025), we analyzed 4,245 *TP53* mutation events in 10,967 unique tumor samples from 32 cancer studies, covering 27 cancer types. Mutations were classified into missense, nonsense, frameshift, in-frame indels, splice-site variants, and fusions, and cancer types were grouped for comparative analysis. Missense mutations were most common, followed by nonsense and frameshift variants. Female reproductive and gastrointestinal cancers had the highest mutation burdens, while central nervous system tumors showed a higher proportion of missense events. These findings highlight both the ubiquity and cancer type–specific patterns of *TP53* alterations, providing a reference for future functional and clinical studies.

## 2. Introduction

The *TP53* gene, often called the “guardian of the genome,” is a tumor suppressor gene that helps prevent cancer by controlling cell division, repairing DNA damage, and triggering cell death when needed. [1] It is located on chromosome 17 and is one of the most frequently mutated genes in many cancer types, contributing to tumor development and progression. Mutation in TP53 can disable these protective functions, allowing cells to grow uncontrollably and form tumors. Different mutation types, such as missense, nonsense, frameshift, and splice site mutations, can variably affect the gene’s function. [2], [3] Moreover, the distribution and prevalence of these mutation types can differ across cancer types due to distinct biological contexts and mutational processes. Understanding the mutational landscape of *TP53* can provide insights into cancer biology and potentially inform therapeutic strategies.

This study aims to characterize the mutational landscape of *TP53* by analyzing the distribution of mutation types across various grouped cancer types. Using publicly available cancer genomics data, the study seeks to identify patterns and differences in *TP53* mutation profiles, enhancing our understanding of its role in diverse cancers.

## 3. Methods

This study employed a bioinformatics approach to analyze *TP53* mutation profiles across multiple cancer types using publicly available genomic datasets. The methodology was designed to ensure reproducibility and clarity, encompassing four main stages: data retrieval, preprocessing, cancer type grouping, and statistical visualization. Mutation data were sourced from the TCGA Pan-Cancer Atlas via the *cBioPortal* platform, curated for accuracy, and systematically categorized into defined mutation classes. To facilitate meaningful interpretation, cancer types were consolidated into broader groups, and descriptive statistics were applied to determine the distribution and frequency of mutation types. Visual representations were generated to highlight key trends and inter-group differences.

### 3.1 Data Source

Mutation data for the *TP53* gene were obtained from *cBioPortal* [4] for Cancer Genomics (https://www.cbioportal.org/) by querying the TCGA Pan-Cancer Atlas studies. On 9 August 2025, *TP53* was searched in the TCGA section, and the “Mutations” tab was selected to retrieve all available somatic mutation data. The complete output, including study identifiers, cancer type annotations, and mutation details, was downloaded in tab-separated values (TSV) format directly from *cBioPortal*.

### 3.2 Data Preprocessing

The raw data were imported into Microsoft Excel for preprocessing. Non-essential columns were removed, and the dataset was reformatted for clarity. Multiple mutation records corresponding to the same sample were merged. Cancer type labels were consolidated into broader categories to simplify analysis. Mutation events were classified into six categories: missense, nonsense, frameshift (insertions and deletions combined), in-frame (insertions and deletions combined), splice-site (including splice region) mutations, and gene fusions.

### 3.3 Grouping of Cancer Types

To improve clarity in visualization and interpretation, the 27 cancer types were grouped into ten broader categories based on tissue origin and biological similarity. These groups include Genitourinary Cancers, Gastrointestinal Cancers, Female Reproductive Cancers, Central Nervous System Cancers, Hematologic Cancers, Skin & Melanocytic Cancers, Endocrine Cancers, Thoracic Cancers, Head & Neck Cancers, and Sarcomas. The full mapping of original cancer types to grouped categories, along with the corresponding number of *TP53* mutation events, is provided in Table 1.

**Table 1.** Distribution of TP53 Mutations Across Cancer Categories. The table presents the number of TP53 mutations identified in various grouped cancer categories. Each category includes specific cancer types as classified in cBioPortal. The data highlight variation in TP53 mutation frequency across cancer types, with the highest counts observed in female reproductive cancers and gastrointestinal cancers.

| Grouped Cancer Category | Original Cancer Type(s) from cBioPortal | TP53 Mutation (s) |
| --- | --- | --- |
| Central Nervous System (CNS) Cancers | Glioblastoma, Glioma | 469 |
| Endocrine & Neuroendocrine Cancers | Adrenocortical Carcinoma, Pheochromocytoma, Thyroid Cancer | 23 |
| Female Reproductive Cancers | Breast Cancer, Cervical Cancer, Endometrial Cancer, Ovarian Epithelial Tumor | 1039 |
| Gastrointestinal Cancers | Cholangiocarcinoma, Colorectal Cancer, Esophagogastric Cancer, Hepatobiliary Cancer, Pancreatic Cancer | 963 |
| Genitourinary Cancers | Bladder Cancer, Prostate Cancer, Renal Clear Cell Carcinoma, Renal Non-Clear Cell Carcinoma, Seminoma | 346 |
| Head and Neck Cancer | Head and Neck Cancer | 436 |
| Hematologic Cancers | Leukemia, Mature B-Cell Neoplasms | 25 |
| Sarcoma | Sarcoma | 99 |
| Skin & Melanocytic Cancers | Melanoma | 83 |
| Thoracic Cancers | Non-Small Cell Lung Cancer, Pleural Mesothelioma, Thymic Epithelial Tumor | 762 |

### 3.4 Data Analysis and Visualization

Mutation counts were aggregated by mutation type within each cancer group. Frequencies and percentages were calculated for both overall and group-specific distributions. Data visualization was performed using Microsoft Excel and online charting tools to generate two primary figures: one summarizing the overall mutation type distribution and another showing mutation frequencies across the defined cancer groups.

## 4. Results

Mutation data for *TP53* were obtained from a combined dataset of 10,967 tumor samples across 32 studies available on *cBioPortal*. In total, 4,245 *TP53* mutations were identified and analyzed, classified by mutation type and cancer group. The analysis covered 27 cancer types grouped into ten biologically relevant categories.

### 4.1 Overall Mutation Type Distribution

Overall, missense mutations were the most frequent, accounting for 64.3% (n = 2,731) of all *TP53* mutations, followed by nonsense mutations (12.9%, n = 546), frameshift mutations (12.3%, n = 522), splice-site mutations (7.9%, n = 335), in-frame mutations (2.1%, n = 91), and fusion events (0.5%, n = 20).

**Figure 1.**
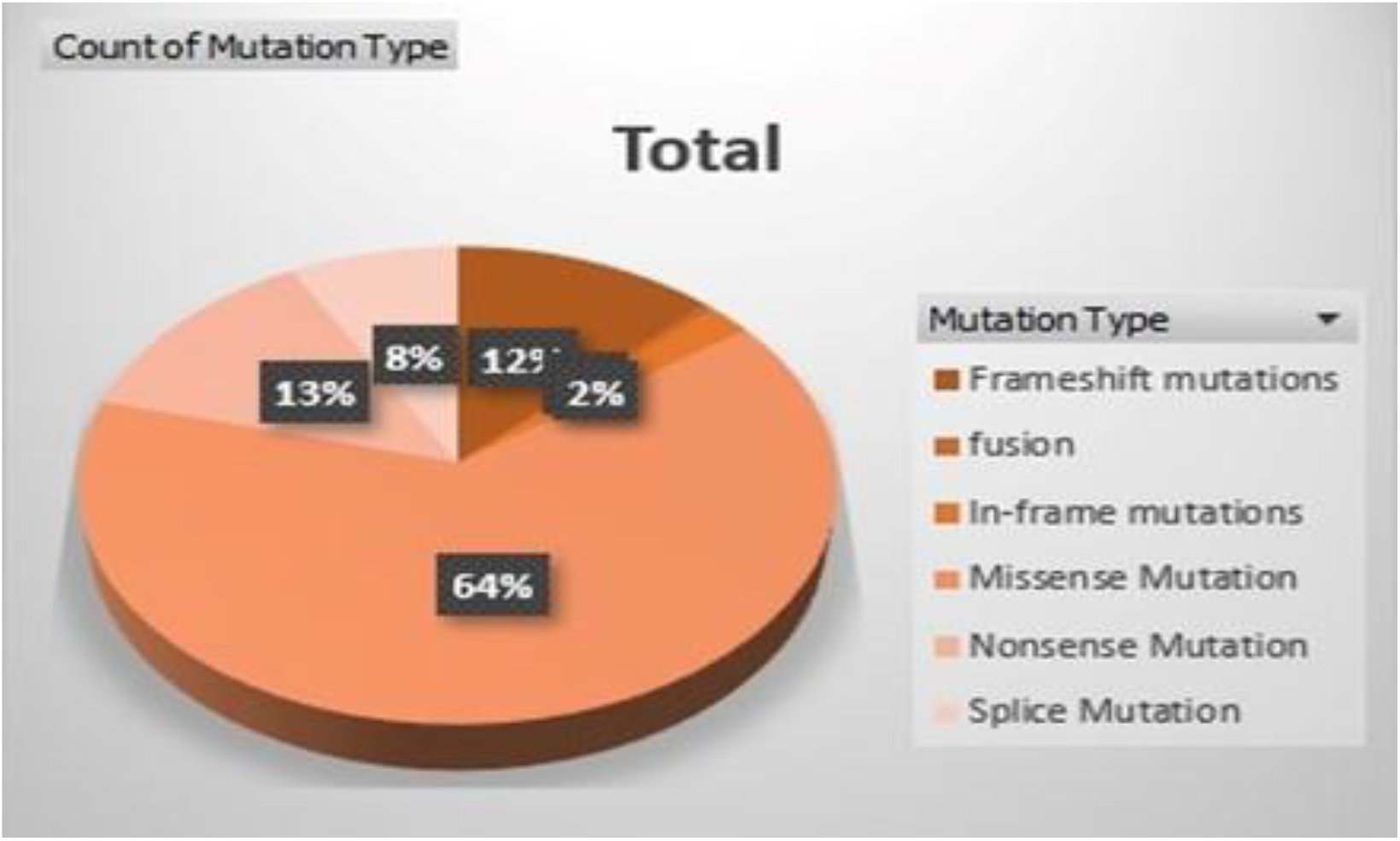
Pie chart showing the distribution of 4,245 TP53 mutation types across 10,967 tumor samples from 32 studies. Missense mutations dominate, followed by nonsense, frameshift, splice-site, in-frame mutations, and fusion events.

### 4.2 Mutation Distribution by Cancer Group

The mutation landscape of *TP53* varied across cancer groups. Female reproductive cancers showed the highest overall burden (n = 1,039), predominantly missense (64.4%, n = 669), followed by frameshift (14.6%, n = 152) and nonsense mutations (11.5%, n = 119). Gastrointestinal cancers (n = 963) displayed a similar missense predominance (63.9%, n = 615) but a comparatively higher frameshift proportion (13.4%, n = 129). Thoracic cancers (n = 762) had 63.2% missense mutations (n = 481) and the highest splice mutation frequency (10.6%, n = 81). In head and neck cancers (n = 436), missense mutations accounted for 57.8% (n = 252), with frameshift mutations representing 14.7% (n = 64). Genitourinary cancers (n = 346) had a slightly lower missense share (67.9%, n = 235) and a balanced distribution of other mutation types. CNS cancers (n = 469) had the highest proportion of missense mutations among all groups (76.5%, n = 359), despite a moderate mutation count. Hematologic cancers exhibited the fewest TP53 mutations overall (n = 25), with 56.0% missense (n = 14) and 16.0% frameshift variants (n = 4). Sarcoma (n = 99), skin & melanocytic cancers (n = 83), and endocrine & neuroendocrine cancers (n = 23) presented smaller mutation counts but retained the missense-dominant pattern (51.5%, n = 51; 56.6%, n = 47; and 34.8%, n = 8, respectively).

**Figure 2:**
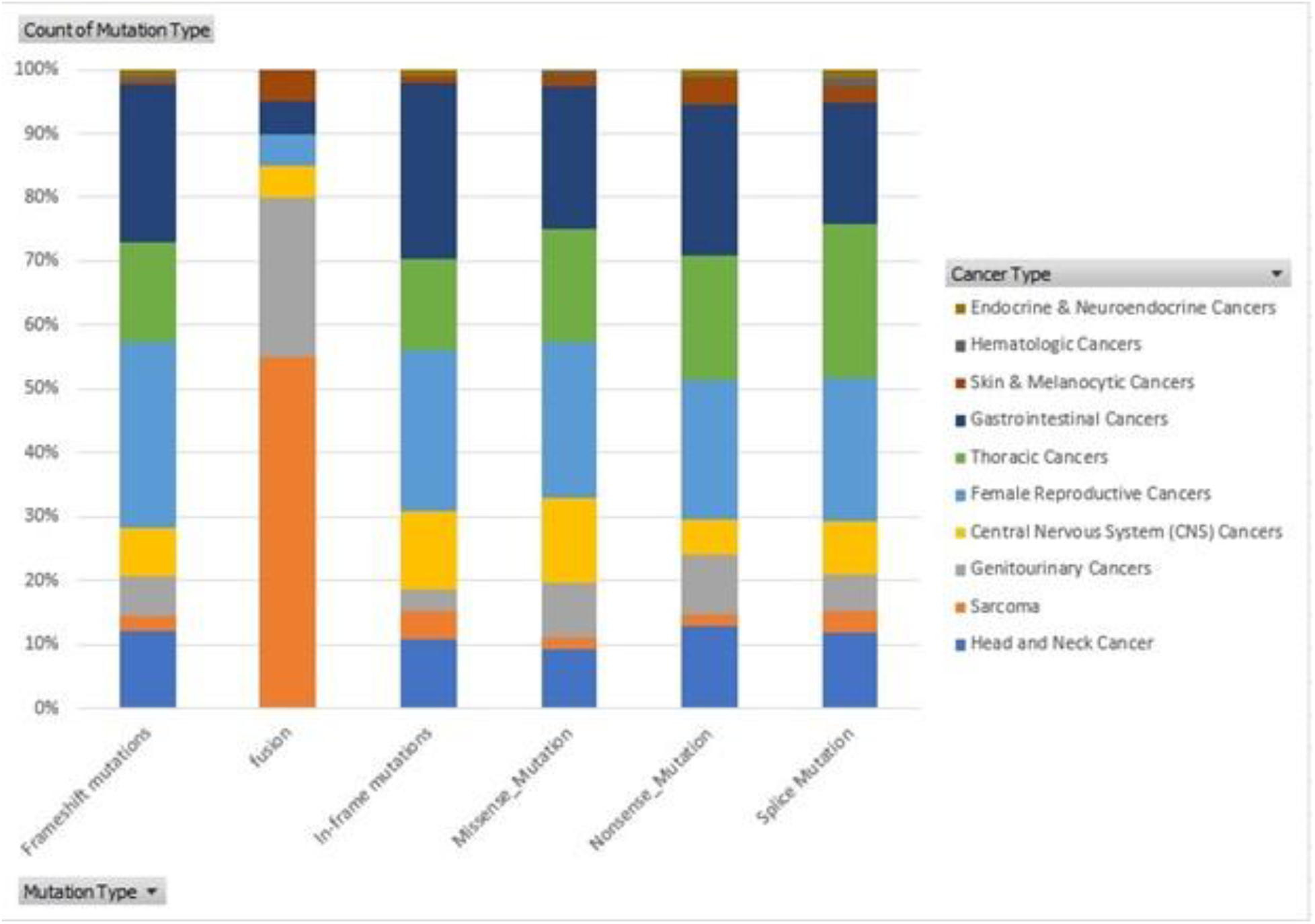
Proportional distribution of cancer types across six mutation categories: Frame Shift, Fusion, Inframe, Missense, Nonsense, and Splice mutations. Each stacked bar shows the percentage contribution of different cancer types, color-coded for Endocrine & Neuroendocrine, Hematologic, Skin & Melanocytic, Gastrointestinal, Thoracic, Female Reproductive, Central Nervous System (CNS), Genitourinary, Sarcoma, and Head & Neck cancers. The chart illustrates variation in cancer prevalence by mutation profile.

### 4.3 Notable Observations

*TP53* mutations were predominantly missense across all cancer groups, peaking in central nervous system cancers (76.5%). Female reproductive cancers had the highest mutation burden with notable frameshift (14.6%) and nonsense (11.5%) rates. Gastrointestinal cancers also showed elevated frameshift mutations (13.4%), while thoracic cancers had the most splice-site mutations (10.6%). Hematologic cancers had the fewest *TP53* mutations, suggesting rarity in this group. Smaller cancer types followed similar missense-dominant patterns.

## 5. Discussion

In this multi-study analysis of 10,967 tumor samples from 32 cancer studies, *TP53* mutations were found to be widely distributed across multiple cancer types, with marked variability in frequency. The highest mutation prevalence was observed in head and neck cancer, non-small cell lung cancer, and esophagogastric cancer, whereas certain malignancies such as leukemia and adrenocortical carcinoma showed relatively low mutation rates. These findings are consistent with the established role of *TP53* as one of the most frequently mutated tumor suppressor genes across human cancers, particularly in malignancies linked to environmental carcinogens or genomic instability.

Our results broadly align with previous large-scale datasets such as TCGA and ICGC, which also report high *TP53* mutation rates in smoking-associated cancers and tumors with high mutational burden. However, the relatively low mutation frequencies in some cancers may reflect biological differences in tumorigenesis, alternative driver mutations, or the influence of study-specific sampling strategies.

The biological and clinical implications of these patterns are significant. High *TP53* mutation frequencies may contribute to aggressive tumor behavior, resistance to therapy, and poor prognosis.[5] Conversely, tumors with lower *TP53* mutation rates may rely on other oncogenic pathways, which could influence biomarker-driven treatment approaches. [6]Understanding these differences can guide the design of cancer-type– specific therapeutic strategies, including the development of drugs that target p53 restoration or exploit vulnerabilities caused by TP53 loss. [2]

This study benefits from a large, diverse dataset, allowing broad comparisons across cancer types. However, limitations include the absence of detailed clinical outcomes, lack of treatment history, and heterogeneity among the contributing studies. Additionally, this analysis was restricted to mutation frequency and distribution; functional consequences of specific variants were not explored.

Future work could expand this analysis by integrating *TP53* mutation status with clinical data, co-mutation patterns, and pathway enrichment analysis. Such integrative approaches may yield deeper insights into the role of *TP53* mutations in tumor progression and therapy resistance, ultimately aiding precision oncology efforts.

## 6. Conclusion

This analysis highlights the variable distribution of *TP53* mutations across different cancer types, with particularly high frequencies in head and neck, non-small cell lung, and esophagogastric cancers. These patterns reinforce the critical role of *TP53* as a central tumor suppressor and suggest that its mutation burden may influence cancer behavior and therapeutic response. While this study provides valuable insights using publicly available genomic data, further research integrating clinical and functional information is needed to fully understand the prognostic and therapeutic implications of *TP53* mutations.

